# Wound-induced eyespots on butterfly wings at the intersection of immune response and pigmentation development

**DOI:** 10.1101/2025.06.22.660915

**Authors:** Maria Adelina Jerónimo, Ana Rita Garizo, Guilherme W Atencio, David Duneau, Patrícia Beldade

## Abstract

Butterfly eyespots are striking examples of evolutionary novelty arising through the repurposing of ancestral genetic pathways, including pathways involved in wound healing. Given the activation of the immune system during wound healing, and the known links between immunity and pigmentation, we hypothesized that the immune response triggered by wounding contributes to the formation of wound-induced ectopic eyespots on butterfly wings. We tested this hypothesis by wounding the wings of *Bicyclus anynana* pupae and modulating immune challenge levels with application of varying quantities of heat-killed bacteria. These led to up-regulation of defense-related genes in the wounded wing as well as in the contralateral non-wounded wing. Immune activation did not affect the likelihood of ectopic eyespot formation, but it did significantly influence the extent of pigmentation changes around injury sites: stronger immune activation produced larger ectopic eyespots. This effect was localized to the treated wing and did not affect contralateral, control wound-induced eyespots. Additionally, stronger immune challenges led to smaller overall wing size and to relatively smaller native eyespots. Our findings reveal that immune activation contributes to the development of wound-induced eyespots and impacts both pigmentation pattern formation and wing size regulation. These results underscore the complex interplay between immune function and developmental processes, and provide new insights into the origins of lineage-specific morphological innovations.

## INTRODUCTION

Butterfly wing patterns are a prime example of ecologically significant and evolutionarily diversified features that have helped elucidate both proximate (molecular) and ultimate (evolutionary) mechanisms underlying phenotypic diversity (e.g. Nijhout 1991, Beldade and Brakefield 2002, Joron et al. 2006, Sekimura and Nijhout 2017, McMillan et al. 2020, Marcus 2021, Teng and Zhang 2024). Butterfly eyespots, in particular, which are involved in mate choice and predator avoidance, have been models of various research topics in evo-devo (Beldade and Peralta 2017, Beldade and Monteiro 2021), including developmental constraints (e.g. Beldade et al. 2002a, Allen et al. 2008) and modularity (Beldade et al. 2002b, Beldade and Brakefield 2003, Monteiro et al. 2003), developmental plasticity (e.g. Bhardwaj et al. 2020, van Bergen et al. 2024), and evolutionary novelty (e.g. Saenko et al. 2008, Beldade and Saenko 2009). Previous studies have uncovered genetic similarities between the development of the eyespots characteristic of Nymphalid butterflies and various ancestral processes that extend beyond Lepidoptera. These processes include the development of legs (Carroll et al. 1994, Saenko et al. 2011), embryos (Saenko et al. 2010, Tong et al. 2014), and antenna (Murugesan et al. 2022), as well as wound healing (Özsu and Monteiro 2017). This study aims at investigating the connection between the immune system activation that accompanies wound healing and the formation of butterfly eyespots.

Eyespots consist of concentric rings of different colors produced around central organizers (Iwasaki et al. 2017). These organizers are specified in the larval stage and instruct color ring formation during pupal development (reviewed in Beldade and Monteiro 2021). In early pupae, presumptive eyespot centers act as sources of diffusible signaling molecules whose concentration gradients determine the production of different pigments in scales at various distances (Figure 1). In addition to native eyespots, several Nymphalid butterflies are capable of producing ectopic eyespots at the sites of injuries inflicted on early pupal wings (Nijhout 1985, Brakefield and French 1995). These wound-induced ectopic eyespots point to a mechanistic overlap between wound healing and eyespot development (e.g. Saenko et al. 2008, Beldade and Saenko 2009). It has been proposed that wounds mimic eyespot organizers by acting as local sources or sinks of molecular signals that influence the color fate of nearby wings scales (Nijhout 1985, French and Brakefield 1992, Brakefield and French 1995). While wounding has long been used as a tool to investigate properties of the wing epidermis affecting eyespot formation (e.g. Brakefield et al. 2009a, Otaki 2011), the mechanisms and molecular players by which injury sites elicit organized pigmentation patterns remain largely unknown.

**Figure 1.**
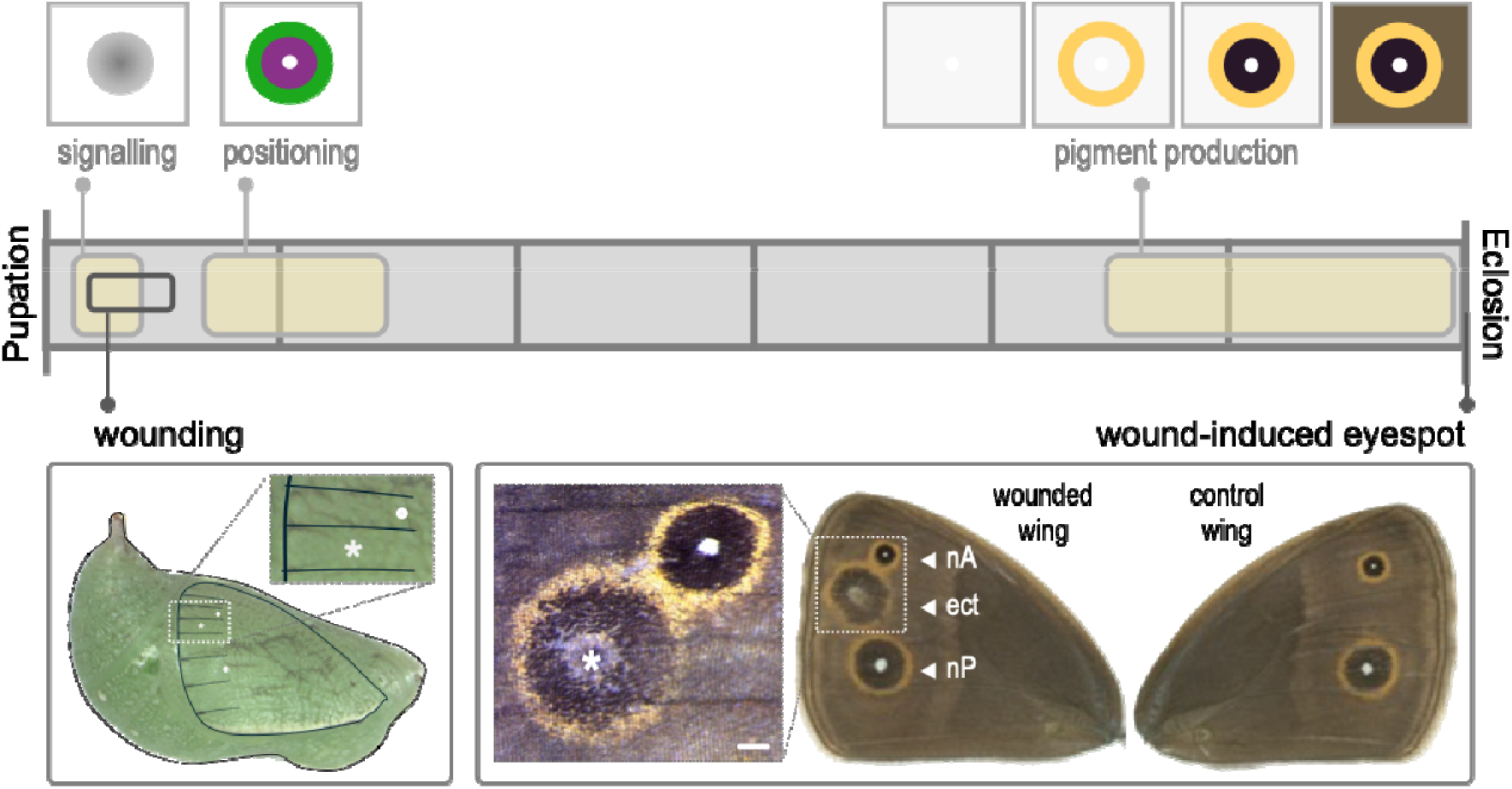
Eyespot formation in Bicyclus anynana pupal wings. Horizontal grey bar in the center represents the duration of pupal life, from pupation until adult eclosion, typically lasting ca. 6 days at 27 °C. Yellow-shaded areas inside the bar represent the approximate timing of the three main phases of native eyespot development that occur during pupal life (schemes above the bar; cf. Beldade and Monteiro 2021): production and diffusion of a morphogen from presumptive eyespot centers (signaling phase), expression of distinct transcription factors defining future color rings (positioning phase), and actual pigment production. Wound-induced ectopic eyespots are illustrated below the bar. Left panel refers to the wounding of the dorsal surface of the forewing through the pupal cuticle, 6-12 hr post-pupation. Photo displays a lateral view of a pupa (head to the right). Superposed to the photo of the green pupa, a drawing outlines the contour of the pupa, its left forewing, and the distal portion of wing veins (black lines), as well as the location of the two presumptive centers of native eyespots (white circles), and of wound sites (white *). Wounded pupal forewing on the left panel is in the same orientation as the corresponding adult forewing on the right panel. Right panel shows native and wound-induced eyespots visible on the dorsal surface of the adult forewings. White arrowheads point at eyespots on wounded forewing: native anterior eyespot (nA), wound-induced ectopic (ect), and native posterior eyespot (nP). Inset displays close up of wing area with nA and wound-induced eyespots, with * marking the wound site (size bar = 1mm).

Wound healing is a well-characterized and evolutionarily conserved process (Moussian and Uv 2005, Krautz et al. 2014, Arenas Gómez et al. 2020). Wounding triggers several genetic cascades that ensure the re-epithelization required to close the wound, and the immune response necessary to fight the infections that often follow open wounds (Galko and Krasnow 2004, Krautz et al. 2014, Raziyeva et al. 2021). However, it remains unclear which components of these two processes contribute to the development of wound-induced ectopic eyespots on butterfly wings. Previous research has documented the expression of eyespot development genes around wound sites (Monteiro et al. 2006) and the expression of wound-healing genes in eyespot organizers (Özsu and Monteiro 2017). A transcriptomic study further identified global changes in gene expression following eyespot-inducing wounds, although it was limited to a single developmental time point (Murugesan et al. 2022). Given the close links between pigmentation and immunity (e.g. Wittkopp and Beldade 2009, Sugumaran and Barek 2016, Whitten and Coates 2017, Koike and Yamasaki 2020), we aimed to investigate the role of immune activation in the development of wound-induced ectopic eyespots.

Using *Bicyclus anynana* butterflies as our experimental model (Brakefield et al. 2009c), we tested the hypothesis that immune activation induced by wounding contributes to the formation of wound-induced ectopic eyespots. First, we showed that wounds on pupal wings alter the expression of several immunity-related genes. We then experimentally modulated immune activation by applying varying quantities of dead bacteria to fresh wounds. While this manipulation did not affect the probability of ectopic eyespot formation, it did influence the size of both wound-induced and native eyespots, as well as overall wing size. We discuss our results in light of the strong connections between immunity and pigmentation development, and how our findings can provide the basis to further studies about the mechanism and evolution of those connections.

## RESULTS

We used experimental populations of *Bicyclus anynana* butterflies to investigate the relationship between immunity and the formation of wound-induced pigmentation patterns. Specifically, we aimed at testing the hypothesis that immune activation contributes to the development of ectopic eyespots following injury. If this hypothesis is correct, we expected: 1) eyespot-inducing wounds to elicit a local immune defense response, and 2) varying levels of immune challenge to affect the ectopic eyespots produced around injury sites.

### Eyespot-inducing wounds activate genes related to immunity and melanogenesis

To establish that eyespot-inducing wounds on the dorsal forewings of *B. anynana* pupae trigger a local defense response, we analyzed changes in the expression of genes with known roles in immune defense, selected as indicators of immune activation (Table S1). We used quantitative-real time PCR (qPCR) to characterize changes in levels of gene expression and *in situ* hybridization (ISH) to characterize spatial patterns of gene expression in wounded pupal forewings (Figure 2), following wounds inflicted at specific developmental stages and locations known to induce ectopic eyespots.

**Figure 2.**
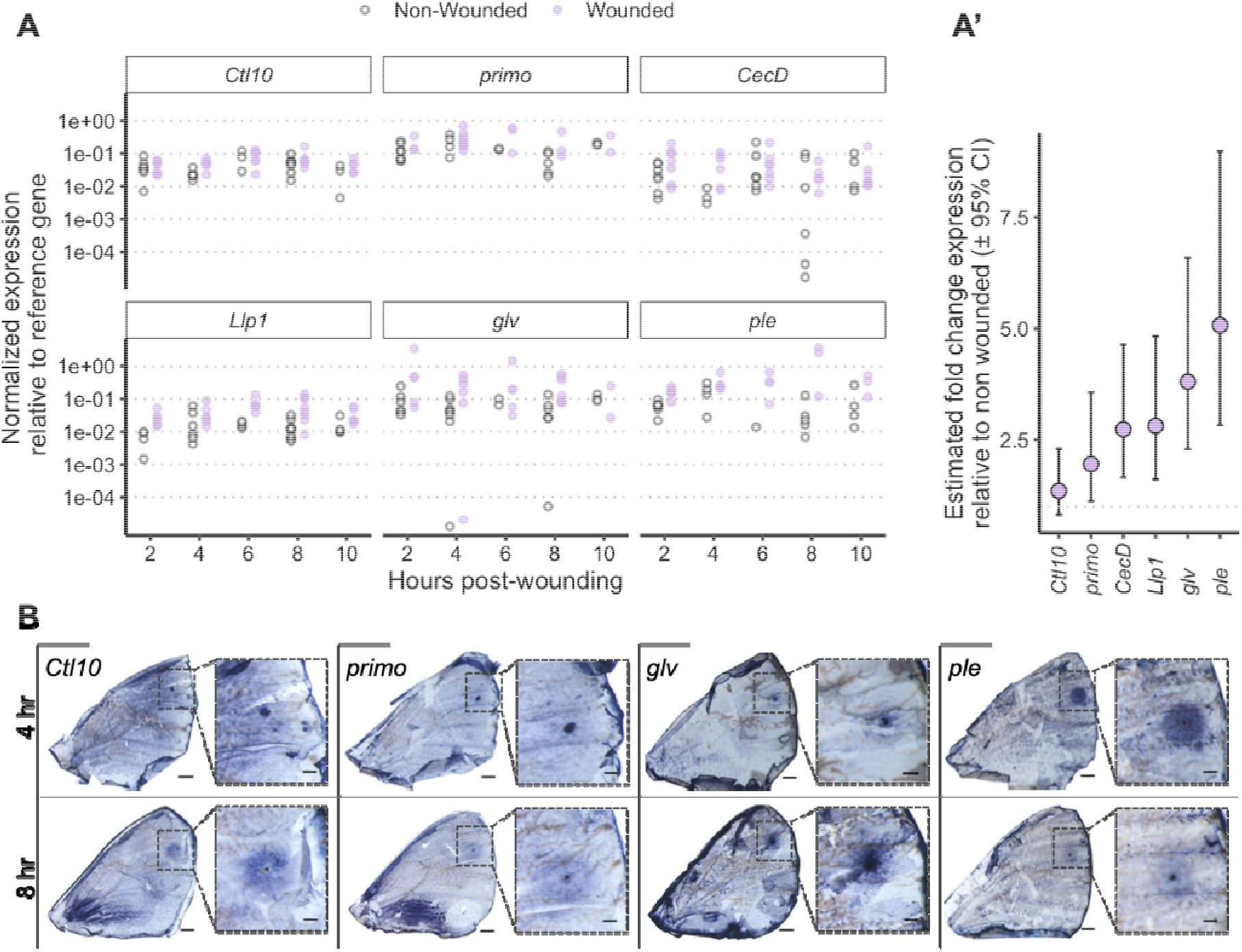
Wound-induced change in expression of genes associated with defense functions. **A**. qPCR analysis of levels of expression of immunity-associated genes in pupal forewings from wounded (purple symbols) and non-wounded (empty symbols) individuals. Up to eight biological replicates (each, a dot on plot) were included per gene and time point. **A**’. Estimated fold change in wounded relative to non-wounded pupae, taking into account all time points and technical replicates (see Material and Methods). 95 % confidence intervals not overlapping with 1 (dotted grey line) confirm that wounding significantly up-regulated all genes, except Ctl10. Code and detailed statistics can be found in supplementary material. **B**. ISH analysis of spatial patterns of gene expression confirmed upregulation of target genes around wound sites. Image of dorsal surface of wounded pupal forewings at 4 hr (top) or 8 hr (bottom post-wounding. Images show whole pupal forewings attached to the pupal cuticle, which is necessary to hold the very fragile pupal wings at that early stage of development and is responsible for the brownish irregular background patterns visible through the wings. Below each image of whole wings, there is a higher magnification image of the wing section around the wound site. Scale bars are 0.5 mm and 0.2 mm for lower and higher magnification, respectively.

We quantified expression changes in six immune-related genes at two-hour intervals from 2 and 10 hr post-wounding. We selected target genes encoding: 1) antimicrobial peptides (AMPs; Yi et al. 2014): cecropin D (*CecD*) and gloverin (*glv*); 2) other peptides involved in recognizing and killing pathogens (Cambi et al. 2005, Gandhe et al. 2007, Chapelle et al. 2009): c-type lectin 10 (*Ctl10*) and lysozyme-like protein 1 (*Llp1*); 3) enzymes involved in processing tyrosine, the basis of melanin synthesis (Futahashi et al. 2022), which contributes to insect immunity (Wittkopp and Beldade 2009): tyrosine hydroxylase or pale (*ple*; Lee et al. 2015) and a tyrosine phosphatase, *primo*. Comparison of forewings between wounded and non-wounded (control) individuals revealed that eyespot-inducing wounds led to significant upregulation of immunity-related genes (Figure 2A, A’).

Additionally, ISH was used to characterize the spatial patterns of expression of some of those in pupal forewings dissected at 4 and 8 hr post-wounding. This revealed mRNA accumulation in circular patterns centered on wound sites, consistent with their upregulation around the injury (Figure 2B).

The activation of the immune response is a common response to wounding, whereby the external protective barrier becomes compromised and triggers organisms to get ready to fight potential infection. In this study, the selected genes served as potential indicators of immune activation, which represent a small subset of what is likely a broader transcriptional response to epidermal damage. While these genes were not selected as candidates for directly regulating eyespot formation, their localized upregulation provides strong evidence that immune pathways are activated in the same spatial context where ectopic eyespots later develop.

### Exposure to different doses of heat-killed bacteria elicits different levels of immune response

Having established that injury elicited a local immune response, we wanted to modulate the strength of this response. Towards this, we applied different doses of heat-killed *Escherichia coli* bacteria to fresh wound sites. Using dead bacteria allowed better control of the timing and level of stimulation of the immune system, without the potentially confounding and detrimental effects of bacteria proliferation inside pupae. To verify that our treatments did correspond to varying levels of immune challenge, we quantified different types of responses typically associated with immune defense (Figure 3): duration of pupal development, pupal mortality, presence of melanotic spots, and expression of immunity genes.

**Figure 3.**
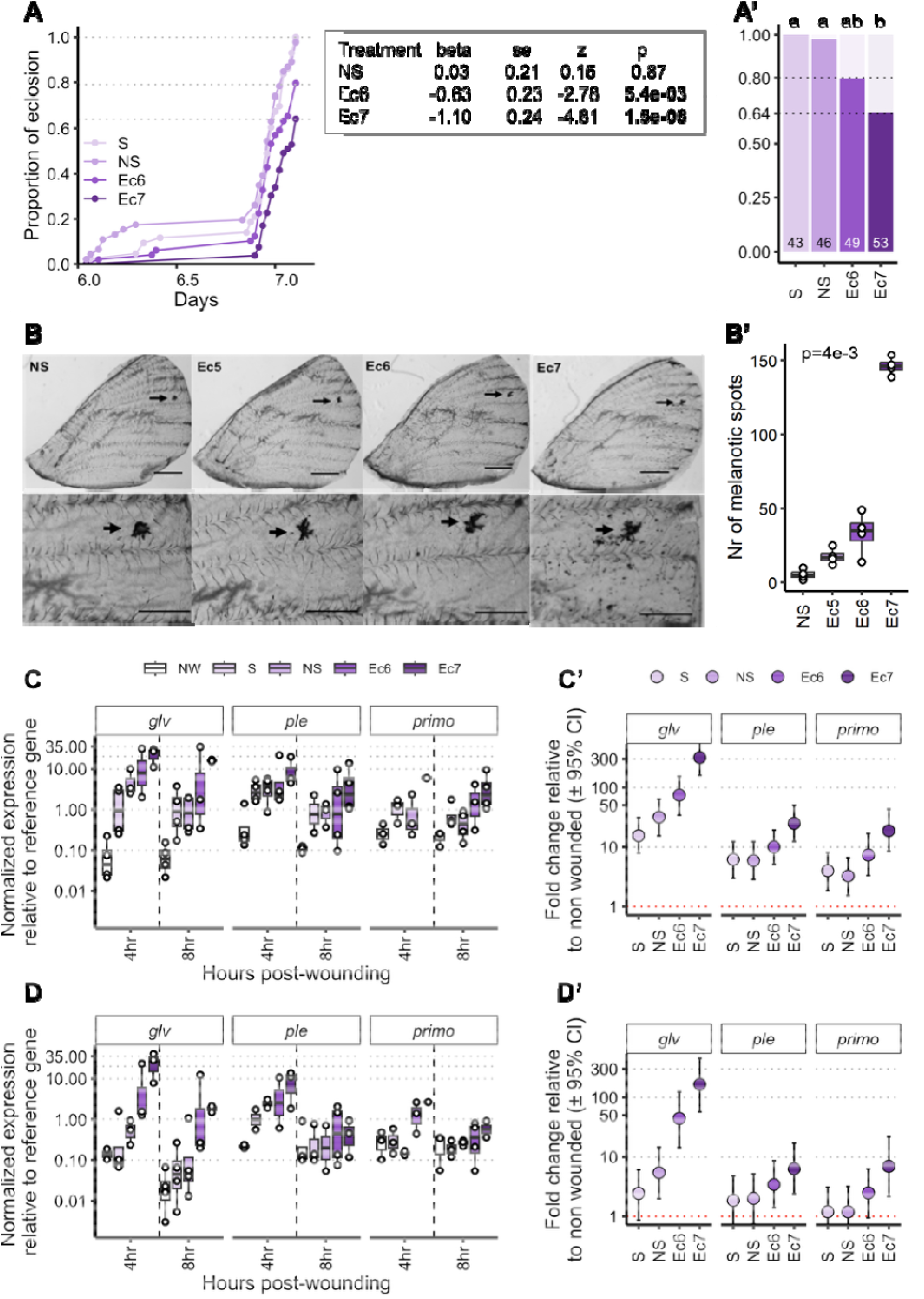
Effects of application of different doses of dead bacteria to eyespot-inducing wounds on traits that reveal differences in level of immune challenge. **A-A’**. Pupal eclosion and survival: higher doses of dead bacteria resulted in slower pupal development (**A**; with the summary of the Cox analysis in the table) and increased pupal mortality (**A**’; with the total number of individuals in each treatment inside the columns). Letters on top of the bars in **A**’ represent the post-hoc comparisons following the binomial linear model used to test the difference in odds of eclosion success. Treatments sharing the same letter are not significantly different. **B-B’**. Melanotic spots: higher concentrations of dead bacteria resulted in a higher number of melanotic spots on the wounded wings. Images (**B**) represent whole dissected pupal wings (top) and zoom of wounded area (bottom). Black dots are the melanotic spots and arrows indicate the wound site. Scale bars are 1 mm and 0.5 mm in whole wings and wing sections, respectively. Plot of counts (**B’**) where each circle refers to counts on one wing. **C-D**. Expression of immunity-related genes: higher concentrations of dead bacteria resulted in increased expression in the wounded pupal wing (**C, C’**) and, to some extent, also on the contralateral non-wounded wing (**D, D’**). In C and D, treatments (X-axis) correspond to non-wounded pupae (NW) and four levels of immune challenge: sterile wounds (S), non-sterile wounds (NS), and application of 10^6^ (Ec6) or 10^7^ (Ec7) dead E. coli to wounds. We studied the expression of three immunity-associated genes (glv, primo, and ple), for both the wounded (**C**) and control (**D**) forewings of up to four replicate individuals (each individual is one circle) dissected at 4 and 8 hr post-wounding. Estimates of expression of each gene relative to non-wounded individuals reported in **C**’ and **D**’ stem from a mixed linear model taking into account all technical replicates and time points. Treatments increased gene expression significantly when the 95 % CI did not overlap with 1 (red dotted line). Code and detailed statistics can be found in supplementary material.

We found that both the duration of pupal development (Figure 3A) and pupal mortality (Figure 3A’; Analysis of deviance for Logistic regression: χ2=31.51, df=3, *p*-value=6.63e-7) differed between treatments varying in the quantity of dead bacteria applied to wounds. While sterile (S) and non-sterile (NS) wounds had no detectable effect on pupation success, application of 10^6^ (Ec6) and 10^7^ (Ec7) dead bacteria resulted in 20 and 36 % mortality, respectively.

We also found that the number of melanotic spots, which are dark pigmented marks often associated with defense responses, differed between treatments (Kruskal-Wallis: χ2=13.26, df=3, *p*-value=0.004), with the highest counts observed in the Ec7 treatment (Figure 3B).

In addition, qPCR analysis of pupal wings revealed that wounds treated with higher bacterial loads exhibited increased expression of the immune-related genes *glv, ple*, and *primo (*Figure 3C-C’). Notably, this upregulation was detected not only in the wounded forewings receiving bacterial application, but also in the contralateral, non-wounded wings of the same individuals (Figure 3D-D’), indicating a systemic immune response to localized immune challenge.

### Immune challenge affects the size but not the likelihood of ectopic eyespot formation

Having established that our treatments elicited different levels of immune response, we next assessed their effects on wound-induced ectopic eyespot formation. We analyzed the wings of the adults eclosed from wounded pupae to assess the effect of immune challenge on the likelihood and size of the ectopic color change (Figure 4).

**Figure 4.**
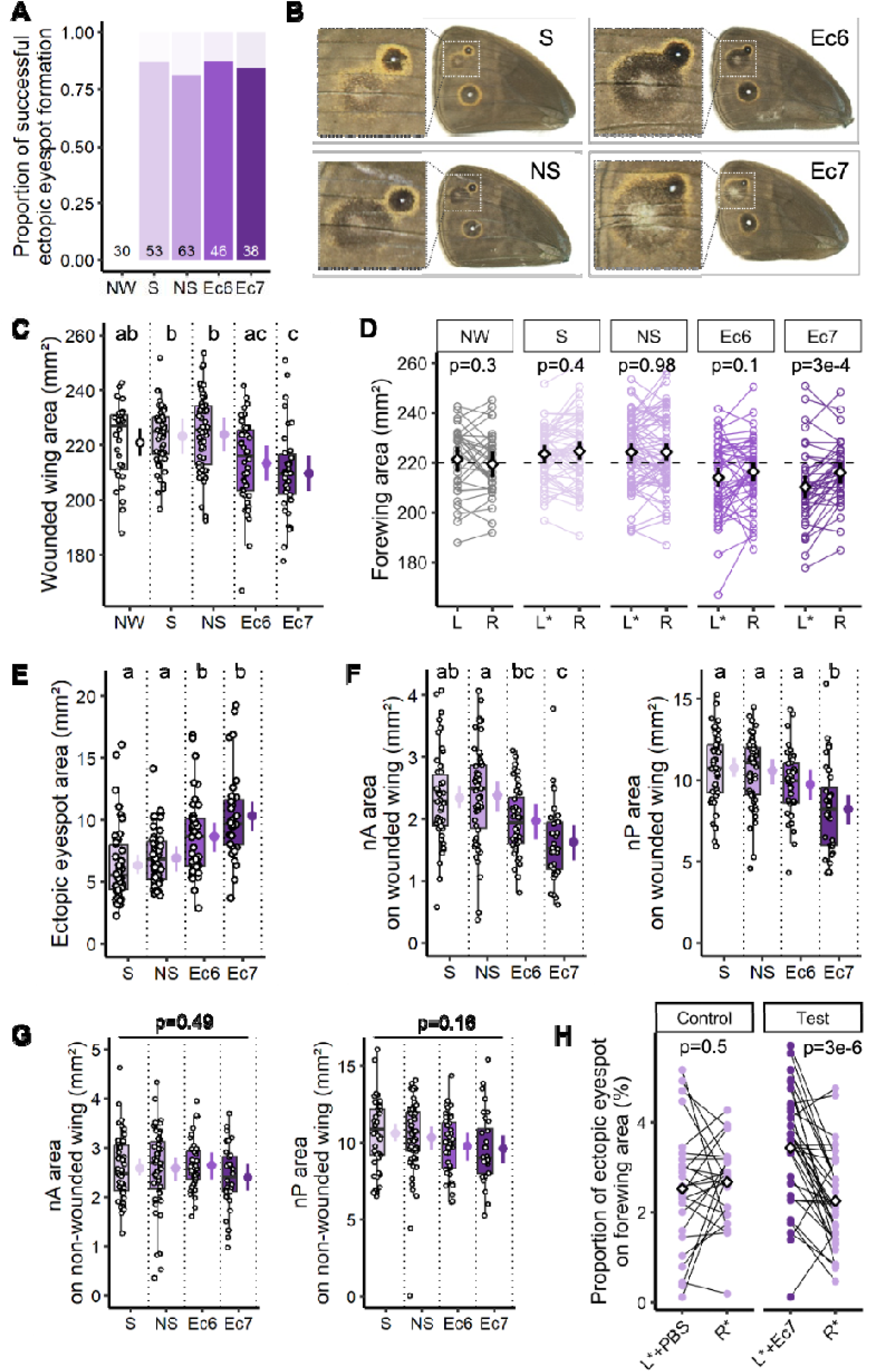
Effects of immune challenge on wing pattern development. Treatments correspond to non-wounded pupae (NW) and different levels of immune challenge applied to forewing wounds: S (sterile wounds), NS (non-sterile wounds), Ec6 (addition of 10^6^ dead E. coli), and Ec7 (addition of 10^7^ dead E. coli). **A**. No effect of immune challenge on the probability of wounds inducing the formation of an ectopic pigmentation pattern. For each bar of immune challenge treatments, stronger color represents individuals that showed an ectopic color alteration. **B**. Images of dorsal surface of wounded forewings of adults from different treatments, with inset highlighting ectopic eyespot near native anterior eyespot. **C-D**. Effect of treatment on wing area: the immune challenge decreased the size of the wounded wing (**C**), which became significantly smaller than the contralateral non-wounded wing for treatment Ec7 (**D**; with the dashed line around mean value for NW treatment). **E**. Effect of treatment on the size of wound-induced ectopic eyespots: the size of the ectopic eyespot increased as the immune challenge increased. **F**. Effect of treatment on native eyespots on the wounded wing: the size of the anterior and posterior eyespots was reduced when the immune challenge applied to the wound increased. This result remained when controlling for the effect of immune challenge on wing size (see text). **G**. No effect of treatment on native eyespots on the contralateral, non-wounded wing: The size of the native eyespots on the contralateral non-wounded wing was not noticeably changed by the treatments. **H**. Effect of immune challenge applied to one wing on wound-induced eyespots on both wings: In all individuals, both wings were wounded, but the wound on the left wing was either exposed to PBS (Control group; corresponding to NS treatment) or to Ec7 (Test group). In all plots, each dot corresponds to measurements of one individual. In **C** and **E-G**, the box plots represent interquartile and median, with whiskers extending to the most extreme data points that fall within 1.5 times the interquartile. To the right of each boxplot, the circle represents mean value and lines represent the 95 % confidence intervals. Letters above each treatment reflect results of post-hoc pairwise comparisons run when treatment had a significant effect: same letter when there is no statistically significant difference (alpha=0.05). In **D** and **H**, fine lines connect measurements for the left (L) and right (R) wings of the same individual, * indicates wounding, and the p-value refers to comparison between left and right. Code and detailed statistics can be found in supplementary material.

Ectopic eyespots were formed in over 81% of cases, regardless of treatment. The probability of ectopic eyespot formation did not vary significantly with the level of immune challenge (Analysis of deviance for logistic regression: χ2=1.02, df=3, *p-*value=0.797; Figure 4A). On the other hand, the size of the wound-induced ectopic eyespots was significantly affected (F-test for linear model: F= 15.043, df=3, *p-* value=1.06e-8): stronger challenge resulted in greater extension of ectopic color change around the wound site (Figure 4B, 4E).

The level of immune challenge also affected wing size, as well as the relative size of native eyespots. Compared to non-wounded individuals, individuals receiving dead bacteria showed a reduction in overall wing size (F-test for linear model: F=9.24, df=4, *p-*value=6.23e-7, Figure 4C), suggesting a cost of the immune response. Within individuals, the difference in size between the wounded and contralateral non-wounded wings of the same individual (i.e. the degree of allometry) was particularly strong as the immune challenge increased (Figure 4D), suggesting growth disruption in response to injury and immune activation.

In addition to these effects, we found that native eyespot size was also influenced by the wound treatments (Figure 4F-G). Both, the anterior native eyespot (nA in Figure 1B; F-test for linear model: F=9.69, df=3, *p-*value=6.34e-6), located adjacent to the wounded compartment, and the posterior eyespot (nP in Figure 1B; F-test for linear model: F=9.55, df=3, *p-*value=7.50e-6), located further away, were reduced in size under stronger immune challenge. Importantly, these effects remained significant even when controlling for the reduction in overall wing size (F-test for linear model: nA: F=6.32, df=3, *p-* value=0.0004; nP: F=8.56, df=3, *p-*value=2.56e-5), indicating that the reduction in native eyespot size was not merely a consequence of allometric scaling, but reflected a specific impact of immune challenge on eyespot development. On the other hand, the native eyespots on the contralateral wings were not noticeably affected (Figure 4G; F-test for linear model: nA: F=0.797, df=3, *p-*value=0.497; nP: F=1.7^2^, df=3, *p=*0.164).

### Local immune challenge triggers systemic gene expression but affects ectopic eyespot size only locally

As previously observed, and consistent with the systemic nature of the immune response, applying heat-killed bacteria to a wound on one forewing triggered changes in gene expression not only at the wound site but also in the contralateral, non-wounded forewing (Figure 3D). This raised the question of whether the effects of immune activation ectopic eyespot size were similarly systemic, or restricted to the site of bacterial application.

To test this, we conducted an experiment in which both pupal forewings were wounded. In test individuals, dead bacteria (107 cells; Ec7) were applied to only one of the wounds, while in control individuals, no bacteria were applied to either wing. Comparing the wound-induced eyespots on the two wings of test individuals, we could determine whether immune challenge affected ectopic eyespot size systemically (both wings should show enlarged eyespots relative to controls) or locally (only the treated wing should show enlargement) (Figure 4H).

Consistent with a localized effect, we found that application of dead bacteria increased ectopic eyespot size on the treated wing but not on the contralateral wounded wing (F-test on linear mixed model: F=13.69, df=2, p-value=1.55e-5 for wing-treatment interaction). The size of ectopic eyespots on wounds with no bacteria did not differ between test *versus c*ontrol individuals; (F-test on linear model: F=1.21, df=2, *p-*value=0.3). Thus, while expression of immunity-related genes responded systemically to localized challenge, the enhancement of ectopic pigmentation remains a localized phenomenon, likely dependent on wound-specific signaling dynamics.

## DISCUSSION

The close interplay between immunity and pigmentation (e.g. Wittkopp and Beldade 2009, Sugumaran and Barek 2016, Whitten and Coates 2017, Koike and Yamasaki 2020), as well as the known activation of the immune system during wound healing (e.g. Krautz et al. 2014), led us to hypothesize that wound-induced immune responses contribute to the formation of ectopic eyespots on butterfly wings. To test this, we used *Bicyclus anynana* as our experimental model (Figure 1), a species well characterized with respect to both the cellular and genetic bases of eyespot formation and variation (reviewed in Beldade and Monteiro 2021), as well as in studies of wound-induced eyespot formation. Prior work has established the spatial and temporal parameters under which wounds lead to ectopic eyespot formation in this species (Brakefield and French 1995), and wounding has been used to interrogate the properties of developing wing tissue during eyespot specification (Brakefield et al. 2009a).

### Summary and interpretation of findings

Our central prediction was that wound-induced ectopic eyespots would vary depending on the level of immune activation associated with the wound. We first confirmed that wounds capable of inducing ectopic eyespots elicit a defense response, as indicated by the upregulation of immunity-related genes (Figure 2). Because complete suppression of the immune response is neither technically feasible nor biologically desirable, we instead modulated immune activation levels by applying different concentrations of heat-killed *E. coli* to fresh wounds. These treatments produced a gradient of immune challenge, as confirmed by gene expression and physiological readouts (Figure 3).

This immune activation gradient had no effect on the likelihood of ectopic eyespot formation (Figure 4A), but it significantly impacted the size of the ectopic color pattern around wounds: stronger immune challenges led to larger ectopic pigmentation areas (Figure 4B, E). This suggests that immune activation influences the *extent*, but not the *initiation*, of the eyespot-inducing signal, potentially by modulating the spatial reach or duration of release of that injury-derived signal.

Interestingly, while the application of dead bacteria to a forewing wound affected immunity gene expression also in the contralateral wing (Figure 3D), this did not translate into larger ectopic eyespots on contralateral wounds (Figure 4H). This discrepancy may result from either a weaker immune activation in the contralateral wing or the absence of locally restricted factors required for ectopic pattern induction. In contrast, we observed that immune challenge reduced wing size on both wounded and contralateral wings, albeit to a smaller extent for the later (Figure 4C, D), pointing to systemic physiological costs or inter-wing developmental coordination. Whether this bilateral size reduction reflects direct effects of circulating immune factors, energetic trade-offs, or developmental compensation remains an open question.

### Mechanistic uncertainty and candidate processes

The specific components of the insect immune response that influence the development of wound-induced eyespots remain unidentified. In this study, we used the expression levels of defense-related genes such as *glv* and *ple* as markers of immune activation, but neither their known functions nor our data suggest a direct role in ectopic eyespot formation. The precise mechanisms by which immune challenge, and wound response more broadly, affect eyespot development remain to be elucidated.

One candidate is the c-Jun N-terminal kinase (JNK) signaling pathway, which has been implicated in regulating organ size (Willsey et al. 2016), wound healing (Bosch et al. 2005, Lee et al. 2017), and immune activation (Tafesh-Edwards and Eleftherianos 2020). JNK signaling also modulates the expression of Wingless and Decapentaplegic (Iida et al. 2019)), two diffusible molecules proposed as morphogens involved in native eyespot patterning (reviewed in (Beldade and Monteiro 2021). However, both wound healing and immune responses involve numerous molecular and cellular players (Lee and Miura 2014, Tsai et al. 2018, Yu et al. 2020), any of which might influence pigmentation development directly or indirectly. These include chemical signals (e.g., wounds releasing an eyespot-inducing morphogen, or similar substances such as calcium; Ohno and Otaki 2015), cellular components (e.g., immune cell recruitment, changes in cellular permeability), and physical factors (e.g., mechanical distortion of the epithelium, or changes in surface tension; Otaki 2018, 2020, 202, Tetley et al. 2019, Nakazato and Otaki 2024). Given this complexity, single-gene perturbations, such as CRISPR-Cas9 knockouts, seem poorly suited to dissecting the multifactorial mechanisms by which immune challenge influences ectopic eyespot development.

### Development-immunity pleiotropy and immunity & pleiotropy

That wounds can trigger eyespot-like pigmentation patterns has led to suggestions that wound response pathways may have been co-opted during the evolution of butterfly eyespots (Nijhout 1985, Brakefield and French 1995, Monteiro et al. 2006, Özsu and Monteiro 2017). The origin of lineage-specific or novel traits often involves the redeployment of ancestral genes shared across taxa (True and Carroll 2002). Research across various species made it clear that the development of those “endless forms most beautiful” that captured Darwin’s attention relies heavily on recurrent use of limited building blocks, inspiring terms such as “genetic toolkit” (Carroll 2008) and “hotspot genes” (Stern and Orgogozo 2011).

Despite attempts to categorize genes into discrete functional groups (e.g. developmental *vs* immune), cross-function pleiotropy is widespread. Insect melanogenesis offers a striking example, with enzymes and substrates involved in body pigmentation also playing roles in immune function and behavior (Wittkopp and Beldade 2009). Many genes traditionally classified as “developmental” are involved in immune defense, and *vice-versa* (e.g. Lemaitre et al. 1995, Sluss et al. 1996, Govind 1999, Nunes et al. 2021, Galambos et al. 2024). The Toll pathway, for example, plays dual roles in embryonic axis specification and innate immunity (Valanne et al. 2011), and has been linked to pigmentation patterning in Lepidoptera (e.g. Nishikawa et al. 2013, KonDo et al. 2017). Our findings showing that immune activation influences the extent of pigmentation patterning on butterfly wings add to this body of evidence. More broadly, these results underscore the deeply intertwined nature of developmental and immune processes, and offer insights into the evolutionary co-option of conserved pathways for novel functions, such as the origin of Nymphalid eyespots.

## CONCLUSIONS

Epidermal injury typically initiates a cascade of cellular and molecular processes that promote wound repair (re-epithelialization) and pathogen defense (immune activation). In some organisms, such injuries also trigger localized pigmentation changes, including ectopic rings of color surrounding wound sites. This phenomenon has been observed across diverse taxa, including moths (Sourakov and Shirai 2020) and fish (Ohno and Otaki 2012), but it is most extensively studied in butterflies. In Nymphalid butterflies, such wound-induced patterns have been proposed to reflect the evolutionary co-option of wound-healing pathways in the origin of eyespots (e.g. Saenko et al. 2008, Beldade and Saenko 2009). Despite the conserved nature of wound healing, the mechanisms by which it leads to ectopic eyespot formation remain poorly understood. Given the well-documented links between pigmentation and immunity (e.g. Wittkopp and Beldade 2009, Sugumaran and Barek 2016, Whitten and Coates 2017, Koike and Yamasaki 2020), we hypothesized that immune activation contributes to the development of wound-induced eyespots. Using *B. anynana*, we tested this hypothesis by comparing pigmentation outcomes in pupae subjected to forewing wounds under different levels of immune challenge. We found that stronger immune activation did not alter the likelihood of ectopic eyespot formation, but significantly increased their size. Our findings suggest that immune signaling modulates the extent, rather than the initiation, of wound-induced pigmentation changes. This raises the possibility that similar immune-related mechanisms may underlie wound-induced pigmentation rings in other taxa (e.g. Ohno and Otaki 2012, Sourakov and Shirai 2020). Additionally, we found that stronger immune challenge was associated with smaller forewings and disproportionately smaller native eyespots. Together, our results highlight a functional link between immune activity and both pattern formation and growth regulation in butterfly wings. Future works can build on these findings to dissect the cellular and molecular mechanisms underlying this relationship, and to further explore how shared pathways can contribute to the evolution of novel traits.

## MATERIAL AND METHODS

### Animals

*Bicyclus anynana b*utterflies from a large, outbred, captive population (Brakefield et al. 2009c) were reared in climate-controlled conditions (Sanyo MLR-351H or Aralab FITOCLIMA 10000 EH incubator): 27 °C (+/-0.5 °C) temperature, 65 % (+/-1 %) relative humidity, and 12:12 hr light:dark cycle. Eggs were collected on a young maize plant from cages with approximately 400 adults fed with fresh banana on wet cotton. Between the second and fourth larval instars, larvae fed *ad libitum* on young maize plants were sexed and female larvae were kept in groups of ca. 200 individuals/cage (47.5 x 47.5 x 47.5 cm; BugDorm-44545). Pre-pupae were collected daily and individually transferred into wells of 25-well plates. Pupation time was recorded during the dark hours via time-lapse photography, with a photo taken each 10 min (Canon 1000D digital camera, Hahnel Giga T Pro 2.4GHz wireless timer remote control).

### Wounds and immune challenge treatments

Wounds were inflicted 12 hr post-pupation using a fine tungsten needle (World Precision Instruments; cat. no. 501317), as described in Brakefield et al. 2009b. The dorsal surface of the left forewing was pierced through the pupal case to which the wing is attached at that stage. Injury was inflicted on the vein-bound wing compartment below the anterior-most eyespot, approximately halfway between the wing margin and the location of the eyespots along the proximal-distal axis of the wing (Figure 1).

To obtain different levels of immune challenge, 0.5 µL of suspensions of heat-killed (30 min at 80 °C) *Escherichia coli* (strain DM09-CFP) were applied to the freshly made wound: 10^5^ (Ec5), 10^6^ (Ec6), or 10^7^ (Ec7) bacterial cells/µL in sterilized PBS. As control, we applied 0.5 µL of sterilized PBS to the wounds under sterile (S; pupal cuticle surface-sterilized with 70 % ethanol and wounds with sterile needle under Bunsen burner) and under non-sterile (NS) conditions. Manipulated pupae were returned to 27 °C to continue development until analysis of pupal or adult wings. Once wounded, the decision to add bacteria or not, as well as the quantity of bacteria to apply, was done randomly to avoid block effects. For all but one of the experiments, only one of the two pupal forewings was wounded and the non-wounded contralateral wing was used as control. For the experiment to distinguish between local and systemic effects of immune activation, wounds were inflicted to both forewings of each pupae, but only the left wing was immune-challenged (by application of 0.5 µL of 10^7^ cells/µL heat-killed *E*.*coli (*Ec7) or control PBS, as described above).

### *In situ* hybridization (ISH) on pupal wings

For the ISH analysis, we dissected forewings attached to the overlaying cuticle from pupae, 4 and 8 hr post-wounding. These were fixed and dehydrated as described in Reed et al. 2011, before storage in methanol at –20 °C. After rehydration (Reed et al. 2011), we followed the ISH protocol for wings described in Saenko et al. 2011. Gene-specific primers for five target genes (Table S1) were used to amplify gene fragments inserted in pGMT-easy plasmid vector (Promega) and to synthesize antisense RNA probes. ISH probes were labeled with digoxigenin (DIG) by *in vitro* transcription with T7 or SP6 RNA polymerase (Promega) in a reaction containing DIG-UTP (DIG RNA labeling mix, Roche) and following manufacturer’s instructions. ISH images were acquired under a stereoscope (Leica MZ6 Stereoscope) coupled to a digital camera (Leica DFC420 C Digital Camera SW Kit).

### cDNA from pupal wings

We extracted forewings from pupae (Brakefield et al. 2009a) representing different treatments, at different time points post-wounding (2, 4, 6, 8, and 10 hr), as well as from non-wounded (NW) pupae as controls. Pupal forewings were collected directly into 2 mL Eppendorf microtubes with a 7 mm glass bead and 500 µL TRIzol (Invitrogen), and kept at –20 °C until homogenization in Qiagen’s TissueLyser II (5 min at maximum speed). The homogenate was stored at –80 °C until total RNA isolation using the Direct-zol MiniPrep Kit (ZIMO Research) and following manufacturer’s recommendations. RNA was eluted in RNAse-free water, and checked for yield and purity (A260/A280 ratio of > 1.8) in a NanoDrop, and for integrity in a 1 % agarose electrophoresis gel. We used 400 ng total RNA for cDNA synthesis in a 10 μl volume reaction with 40U M-MLV RNase H Minus reverse transcriptase (Promega) and 0.5 μg oligo(dT) following manufacturer’s instructions. The cDNA product was diluted for a final volume of 50 μL and used as a template in qPCR (see below). Using the same template RNA, we synthesized cDNA at least twice independently to use as technical replication for each biological replicate.

### quantitative real-time PCR (qPCR)

We used 1 µL of cDNA solution (prepared as explained above) as a template in 10 µL qPCR reactions with 5 µL iQTM SYBR Green supermix (BioRad) 3.6 µM of each primer. We selected target genes reported to play roles in insect immune defense (Table S1), Ribosomal-like protein 10A (RpL10A) as reference gene, after confirming its expression stability across treatments (non-wounded and wounded individuals) and wings (left control wing and right wounded) (Vandesompele et al. 2002, Teng et al. 2012). For all genes the primer pairs were designed and validated to meet the following criteria (Taylor et al. 2010): 1) different amplicon sizes for cDNA versus gDNA template, 2) primer efficiency between 90 and 110 %. qPCR was run in a 7500 Real-Time thermal cycler (Applied Biosystems) and included an initial step of 95 °C for 10 min and 40 cycles of 95 °C for 20 sec, 59 °C for 20 sec, and 72 °C for 30 sec. Analysis of melting peaks (Ririe et al. 1997) allowed eliminating samples with primer-dimers and unspecific amplifications (SDS software version 1.4; Applied Biosystems). We confirmed 100 % ± 10 % reaction efficiency for each of the target genes (cf. Taylor et al. 2010). With the Cq values for target and reference genes, we calculated normalized expression based on the delta Cq method, a variation of the Livak method (Livak and Schmittgen 2001): *normalized expression = 2 exp –DeltaCq*, where *DeltaCq = Cq target gene – Cq reference gene*.

### qPCR for wound effects

To quantify effects of wounds on gene expression levels, we used six target immunity genes (*Ctl10, CecD, glv, Llp1, ple, primo;* Table S1) and one reference, “housekeeping” gene (*RpL10A)*, two treatments (wounded and non-wounded pupae, W and NW, respectively), five time points (2, 4, 6, 8, and 10 hr post-wounding), eight biological replicates (left forewing from eight individual pupae), and two technical replicates (two cDNA synthesis reactions from RNA isolated from extracted wings). We discarded biological replicates for which only one technical replicate worked and those for which the two technical replicates were very different (coefficient of variation for normalized expression greater than 30 %). The normalized expression for all biological replicates retained was calculated as the average between the corresponding technical replicates. Expression data were analyzed in the R statistical environment (R Development Core Team 2015), testing a linear model with a F-test which considered technical replicates, all time points and all genes together. The corresponding R syntax is *Delta_Cq ∼ Gene + Treatment*: *Gene + (1*|*TimePoint) + (1*|*SampleID)*. Confidence intervals for relative fold changes were calculated using parametric bootstrap resampling (n = 999 iterations) with the percentile method.

### qPCR for immune challenge effects

To quantify effects of immune challenge treatments on expression, we used three target genes (*glv, ple, primo)* and one reference gene (*RpL10A)*, five treatments (non-wounded pupae, and four levels of immune system activation: S, NS, Ec6, and Ec7 as described above), two time points (4 and 8 hr post-wounding), two forewings (wounded wing and contralateral, control wing) of each of four biological replicate individuals (four different pupae) and three technical replicates (three cDNA synthesis reactions from RNA extracted from each wing). We discarded technical replicates differing more than 1.5 in delta Cq, and averaged delta Cq for the remainder to obtain the delta Cq for each biological sample and gene combination. Fold change was calculated as the normalized expression of each sample divided by the mean normalized expression of the non-wounded samples for the same gene-wing-timepoint combination. Expression data was analyzed in the R statistical environment (R Development Core Team 2015), testing a linear model with a F-test which considers technical replicates, both time points and all genes together. The corresponding R syntax is *Delta_Cq ∼ Gene + Treatment*: *Gene + (1*|*SampleID) + (1*|*TimePoint) + (1*|*SampleID:Replicate)*. Confidence intervals for relative fold changes were calculated using parametric bootstrap resampling (n = 999 iterations) with the percentile method.

### Immune challenge effects on pupae

Wounded pupae subjected to immune challenge treatments were monitored to assess the duration of pupal development, as well as to quantify wound-induced melanotic spots. We tested the effect of immune challenge levels on the duration of pupal stage using a cox survival analysis (function *coxph* in the *Survival R* package), and on pupal survival using a generalized linear model with a binomial distribution. Post hoc comparisons were then done with a Tukey HSD using the function *glht (*Tukey method) from the package *multcomp*.

We dissected wounded forewings from four pupae per treatment, 4 hr post-wounding. Wings were fixed in 9 % formaldehyde for 30 min and mounted in 100 % glycerol before imaging (Zeiss stereoscope Stemi SV6 attached to a camera UEye Cockpit software under standardized light conditions). Images were loaded onto Fiji software (Schindelin et al. 2012) for counting the melanotic spots in a wing area of 1.60 by 1.20 mm, below the wound site. The effect of treatment (fixed factor with 4 levels, NS, Ec5, Ec6, Ec7) on the number of melanotic spots was assessed using a non-parametric Kruskal-Wallis test. For the post-hoc pairwise comparisons, we used a non-parametric Dunn’s test from the *dunn*.*test* package with FDR correction.

### Immune challenge effects on adult wings

Adults eclosed from pupae from different treatments were allowed to fully stretch their wings, and were sacrificed soon after. Both forewings (wounded and contralateral control) were detached from the body and their dorsal surfaces were scanned (Epson Perfection V600 Photo). These images were analyzed in Fiji (Schindelin et al. 2012) to assess the presence of wound-induced ectopic eyespots, and to measure the area of the native dorsal eyespots (nA and nP in Figure 1) and total wing area.

We tested for differences between immune activation levels (*Treatment*, a fixed factor with four levels: S, NS, Ec6, Ec7) in the likelihood of wounds leading to an ectopic eyespot using a logistic regression model (glm with binomial distribution) followed by a likelihood ratio test (LRT). We tested for differences in wing area and eyespot area using linear models followed by a F-test. We log-transformed the dependent variable when that improved model fit significantly. For the analysis of eyespot size, we also fitted a model using the offset function to assess changes in eyespot size relative to changes in wing size. The corresponding R syntax was: *log_ectopic_area ∼ treatment + offset(log_wounded_forewing_area)*. Post-hoc pairwise comparisons were done using a Tukey test (with alpha = 0.05). Confidence intervals were computed as in previous sections.

### Local vs systemic effects of immune activation on ectopic eyespots

To check if the effect of the immune challenge on ectopic eyespot size was only seen locally (i.e. for the wound where bacteria were applied), we had pupae wounded on both forewings but with bacteria applied only to the wound on the left forewing. As a control for left-right consistency, we had pupae where both forewings were wounded but no bacteria were applied. We compared ectopic eyespots across all wings using a mixed linear model followed by F-test. The corresponding R syntax was: *Ectopic_area ∼ Forewing_area + Treatment + Wing_side:Treatment + (1*|*SampleID). T*he random effect *SampleID a*llowed us to compare within each individual. The size of the forewing was used as covariate to control for the change in wing size. The *Wing_side:Treatment i*nteraction term allowed us to test whether the difference in eyespot size between wings depended on the treatment.

All statistical analyses were done in R (4.5) and R studio. The packages and code used will be made available as supplementary material.

---

**Table S1.**
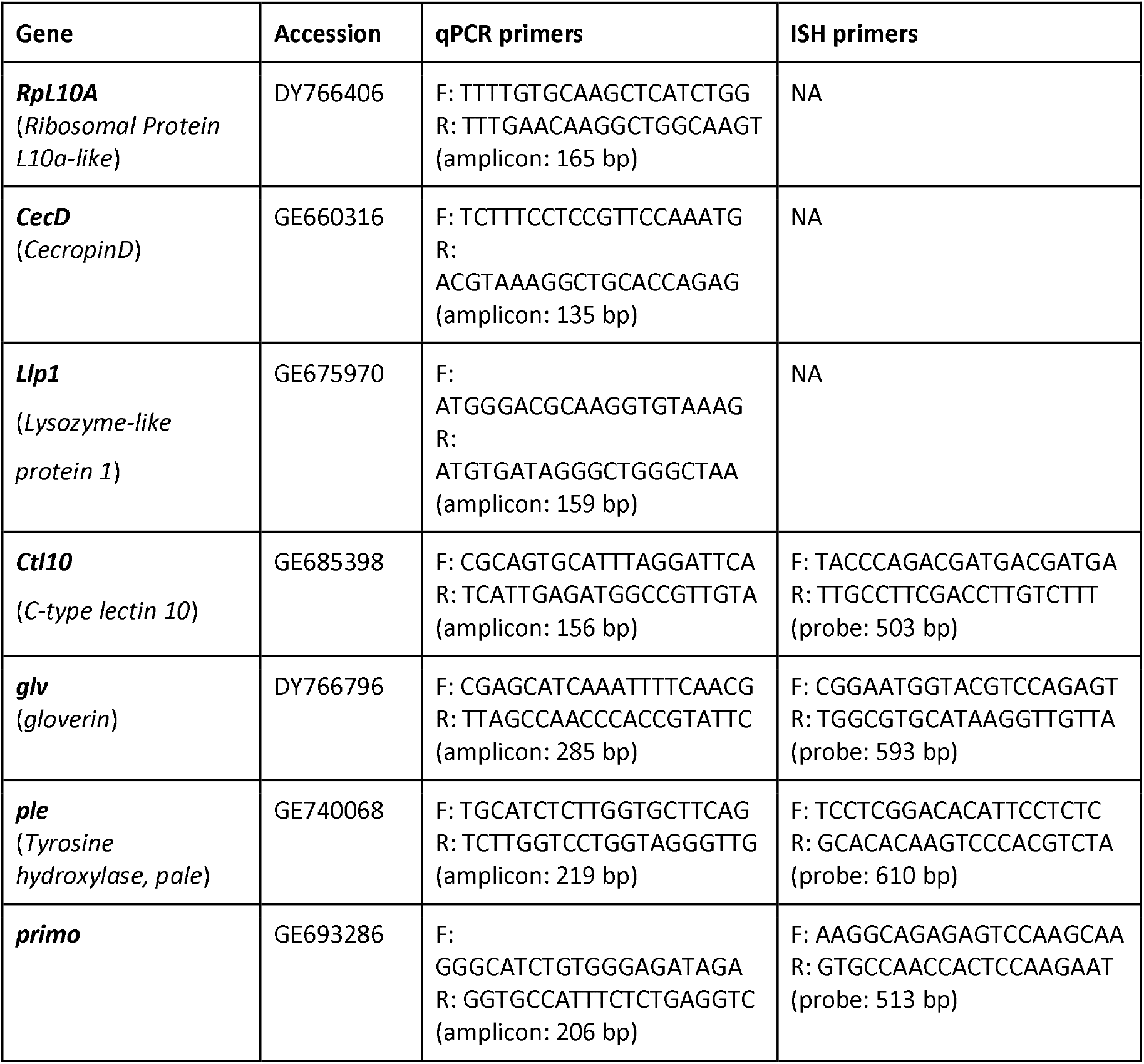
Target genes for expression analysis (qPCR + ISH)

